# A Novel Cell Therapy for COVID-19 and Potential Future Pandemics: Virus Induced Lymphocytes (VIL)

**DOI:** 10.1101/2020.11.26.400390

**Authors:** Rohan Sivapalan, Jinyan Liu, Krishnendu Chakraborty, Elisa Arthofer, Modassir Choudhry, Philip S Barie, Dan H Barouch, Tom Henley

**Author notes:** Co-corresponding Authors: Tom Henley,; Dan Barouch.

## Abstract

The *a priori* T cell repertoire and immune response against SARS-CoV-2 viral antigens may explain the varying clinical course and prognosis of patients having a mild COVID-19 infection as opposed to those developing more fulminant multisystem organ failure and associated mortality. Using a novel SARS-Cov-2-specific artificial antigen presenting cell (aAPC), coupled with a rapid expansion protocol (REP) as practiced in tumor infiltrating lymphocytes (TIL) therapy, we generate an immune catalytic quantity of Virus Induced Lymphocytes (VIL). Using T cell receptor (TCR)-specific aAPCs carrying co-stimulatory molecules and major histocompatibility complex (MHC) class-I immunodominant SARS-CoV-2 peptide-pentamer complexes, we expand virus-specific VIL derived from peripheral blood mononuclear cells (PBMC) of convalescent COVID-19 patients up to 1,000-fold. This is achieved in a clinically relevant 7-day vein-to-vein time-course as a potential adoptive cell therapy (ACT) for COVID-19. We also evaluate this approach for other viral pathogens using Cytomegalovirus (CMV)-specific VIL from donors as a control. Rapidly expanded VIL are enriched in virus antigen-specificity and show an activated, polyfunctional cytokine profile and T effector memory phenotype which may contribute to a robust immune response. Virus-specific T cells can also be delivered allogeneically via MHC-typing and patient human leukocyte antigen (HLA)-matching to provide pragmatic treatment in a large-scale therapeutic setting. These data suggest that VIL may represent a novel therapeutic option that warrants further clinical investigation in the armamentarium against COVID-19 and other possible future pandemics.

## MAIN

Adoptive Cell Therapies (ACT) utilizing autologous patient-derived Tumor Infiltrating Lymphocytes (TIL) have demonstrated reproducible clinical responses in the setting of highly immunogenic tumors such as melanoma and HPV derived cervical cancer. With neoantigen enrichment, this demonstrable clinical efficacy has been extended to common epithelial solid tumors^1, 2^. The efficacy of TIL cell therapy relies on antigen specificity to the cognate cancer neoantigen, and on the ability to expand a large quantity of autologous or allogeneic cancer antigen-specific T cells *ex vivo,* and deliver this number back to the patient to enable cytolysis of the tumor cells and eradication of the cancer^3, 4^. Likewise, antigen-specific T cells are critical for an effective cellular immune response during infection with viral pathogens, and activation of cytotoxic T cells can clear an infection by killing virus-infected cells^5, 6, 7, 8^. Thus, we propose a similar-in-principle novel antigen-specific adoptive T cell therapy for viral infection predicated on precise T cell viral antigen-specificity. However, this viral platform requires cell kinetic expansion in days not weeks, thus allowing for the adoptive transfer of immunologically competent T cells in a therapeutically relevant time course in the setting of acute viral infection.

Like their tumor-resident counterparts, TIL, Virally-Induced Lymphocytes, or VIL, represent those T cells that have been activated in response to TCR-mediated, antigen-specific recognition of protein epitopes from viral particles^9, 10^. Cytotoxic CD8^+^ T cells play a crucial role in mediating viral clearance in response to many respiratory viral infections including respiratory syncytial virus (RSV), influenza and coronavirus (CoV)^5^. Recent evidence has also demonstrated a critical role for the T cell immune response in the pathogenesis of the recently emerged COVID-19 disease, caused by the novel SARS-CoV-2 coronavirus^11, 12, 13, 14^. In addition to the readily detectible humoral immune response in the context of neutralizing antibodies in convalescent patients who had recovered from COVID-19, these studies have collectively shown strong SARS-CoV-2-specific memory T cells are frequently observed^11, 13^. Furthermore, significantly larger T cell responses appear to correlate with severity of the disease, underscoring the importance of T cells above and beyond the humoral antibody response for combating infection^11^. This is an important consideration for a novel therapeutic, as most prophylactic vaccines currently in development for COVID-19 are designed to focus on eliciting antibody responses to the spike protein of SARS-CoV-2^15, 16, 17, 18, 19, 20, 21, 22, 23, 24^.

In addition, the antibody response in recovered COVID-19 patients has been shown to decline several months after infection, raising concerns that therapeutics or vaccines designed to elicit primarily neutralizing antibody responses may not be sufficient to engender the cellular immunity required for long-term duration of protection or to protect from potential repeat infections^25^. Thus, the *a priori* T cell repertoire, both quantity and quality, may portend COVID-19 disease prognosis and may influence the outcome between mild or severe disease. The treatment of patients with convalescent sera has recently been given US FDA Emergency Use Authorization (EUA)^26^. The transfer of neutralizing antibodies reflects a more *passive* serological immune engagement versus the more *active* cellular immune response that SARS-CoV-2-specific T cells would provide.

In an evolution of the cGMP methods for the expansion of TIL for the treatment of cancer that the researchers employ in a currently active human clinical trial^27^, we sought to develop an adoptive T cell therapy for COVID-19 based on rapid *ex vivo* expansion of SARS-CoV-2 antigen-specific VIL. Given the crucial importance of a strong virus-specific T cell response for patients with severe disease, especially during the critical days where respiratory distress is common, the adoptive transfer of a quantity of expanded, activated, effector memory T cells capable of mounting a robust virus-specific response may be important to reduce viral load and improve patient outcomes. Thus, we designed a T cell expansion platform comprising of microbead-based artificial antigen-presenting cells (aAPCs), we termed VIPR-particles (VIL-inducing particles R_x_), that carry MHC pentamer-peptide or tetramer-peptide complexes, specific for immunodominant SARS-CoV-2 epitopes, coupled with costimulatory anti-CD28 antibodies.

Using Peripheral Blood Mononuclear cells (PBMCs) isolated from convalescing COVID-19 patients to represent what may be achievable in the clinic for patients actively suffering from the severe disease, we show that these TCR-specific aAPCs can expand virus-specific VIL up to 1,000-fold over a rapid and minimal culturing duration of just 7-days. Furthermore, these expanded VIL are enriched in virus antigen-specificity, show polyfunctional cytokine responses and acquire a T effector memory phenotype, making them highly suited for participating in an active cellular immune response when adoptively transferred back to patients after this minimal *ex vivo* expansion time. We also demonstrate the broad clinical potential of this platform and its modularity beyond COVID-19 and show in the setting of CMV infection that VIL specific for immunodominant CMV epitopes can also be expanded up to 1,000-fold using CMV-specific VIPR particles over a 7-day culture.

There are unfortunately limited therapeutic options for the treatment of COVID-19 that have demonstrated robust clinical relevance amidst the ongoing pandemic. Considerable, contemporaneous actual experience in the critical care of COVID-19 patients dating from the first quarter of 2020 in New York City, NY, USA, including observations of the clinical time course, hallmark clinical features, and treatment from presentation through critical illness to ultimate resolution or mortality, provides guidance as to the need for a vein-to-vein time of 7-days in order to intercede timely. A protective virus-specific T cell therapy may be an important novel modality for the treatment of SARS-CoV-2 and other pandemic viral pathogens.

## RESULTS

### The detection and enrichment of antigen-specific Virus Induced Lymphocytes (VIL) in CMV infected individuals

To develop a platform for robust and rapid *ex vivo* expansion of viral antigen-specific T cells from patients exposed to viral pathogens, we generated a micro-aAPC capable of providing an immunogenic viral peptide in the context of MHC Class I or MHC Class II molecules in combination with anti-CD28 stimulation molecules (Fig. 1a). MHC-I pentamers or MHC-II tetramers with associated viral antigen peptide complexes, along with anti-CD28 antibodies, were conjugated to 2.8 μm superparamagnetic beads with a monolayer of streptavidin covalently coupled to the surface. We termed these microbead-based aAPCs VIPR particles, and selected immunodominant viral antigens of known MHC specificity to demonstrate their capacity for TCR stimulation and simultaneous expansion of responding T cells. T cells isolated from donor PBMCs were cultured with these VIPR particles for 7-days in the presence of high concentrations of trophic cytokines, IL-2, IL-7 and IL-15 and addition of N-acetylcysteine (NAC), known to improve T cell proliferation^28^. As a validated control to demonstrate efficacy of the platform, we selected an immunodominant MHC-I restricted pp65 antigen of cytomegalovirus (CMV) known to robustly stimulate the TCRs of CMV-specific T cells^29^.

**Fig. 1:**
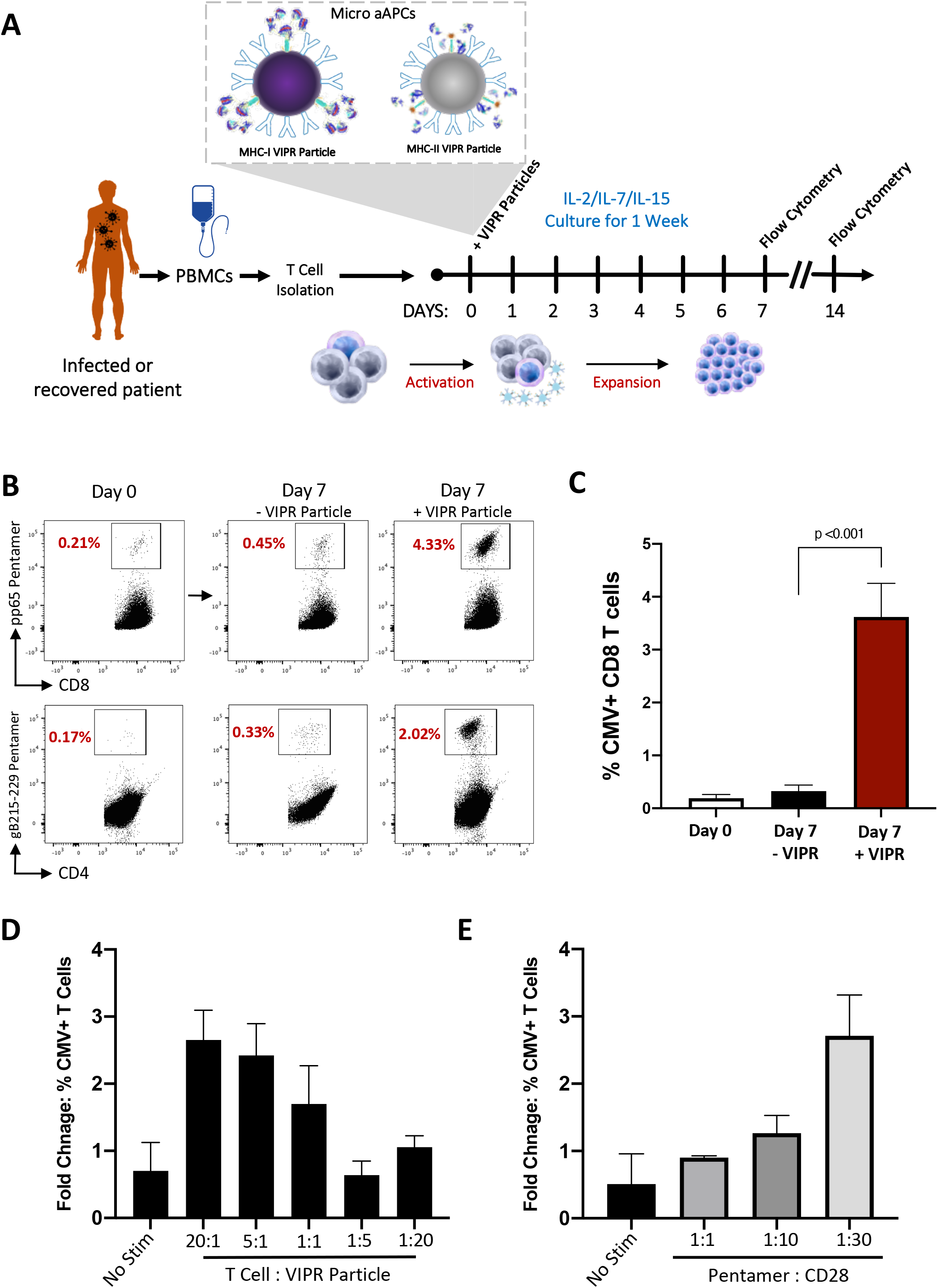
Detection and enrichment of antigen-specific Virus Induced Lymphocytes (VIL) in CMV infected individuals. **a**, Strategy for the isolation, stimulation and enrichment of CMV antigen-specific T cells from donor PBMCs. **b**, Representative flow-cytometric analysis showing proportions of antigen-specific CD8^+^ and CD4^+^ T cells identified by Pentamer staining, 7-days after enrichment and expansion with antigen-specific VIPR Particles. **c**, Summary of CD8^+^ data obtained in b, *(n=3)*. **d**, Histogram plot showing this VIL enrichment is VIPR particles dose-dependent *(n=3)*. **e**, Histogram showing impact on VIL enrichment of MHC-I pentamer and anti-CD28 ratio conjugated to VIPR particles *(n=3)*.

Individuals that have been previously exposed to CMV infection have recirculating virus-specific T cells (VIL) in their blood with TCRs specific for a variety of antigenic CMV peptides^30^. Using fluorescently conjugated pentamers and flow cytometry we analyzed the CD3^+^ T cells isolated from the PBMCs of several independent CMV positive individuals and found approximately 0.2% of T cells were CD8^+^ cells demonstrating specificity for the CMV pp65 antigen (Fig. 1b). After a 7-day culture with VIPR particles, these cells enriched on average 20-fold, reaching Pentamer^+^/CD8^+^ T cell proportions of over 4% (Fig. 1b&c). By comparison, donor T cells cultured in cytokine alone in the absence of VIPR particles showed a minimal enrichment in antigen-specific VIL proportions, even though the T cells proliferated robustly. In addition to the enrichment of virus-specific CD8^+^ T cells using MHC-I restricted antigens, CD4^+^ T cells could also be enriched over 20-fold using MHC-II VIPR particles carrying a well validated CMV glycoprotein antigen (Fig. 1b).

### Optimization of antigen-specific micro aAPCs for T cell expansion

Next we sought to optimize the design of the VIPR particle aAPCs to further enhance the expansion of antigen-specific T cells within the rapid 7-day stimulation culture. To this end, we first investigated the impact that the ratio of T cells to VIPR particles had on the proportion of CMV-specific T cells enriched at day-7. A dose-dependent enrichment was seen with lower doses of particles and higher numbers of T cells, such that an optimal enrichment was observed with a ratio of 20:1 T cells to VIPR particles (Fig. 1d). In addition, we observed that increasing the ratio of molecules of anti-CD28 antibody to peptide-MHC-pentamer also increased the capacity of the VIPR particles for expansion of the antigen-specific VIL population (Fig. 1e)

### The detection and enrichment of SARS-CoV-2-specific T Cells in COVID-19 convalescent individuals

We next evaluated whether VIPR particles can be used to enrich and expand SARS-CoV-2 specific T cells from COVID-19 patients. To investigate this, we obtained PBMCs from COVID-19 convalescent individuals 24 days or more after developing symptoms and testing positive by PCR, and analyzed the frequency and proportion of VIL specific for a recently published and validated immunodominant MHC-I SARS-CoV-2 epitope YLQPRTFLL (YLQ)^31^. Surprisingly, we found YLQ antigen-specific VIL were barely detectible within the isolated T cell populations from these individuals (Fig. 2a&b).

**Fig. 2:**
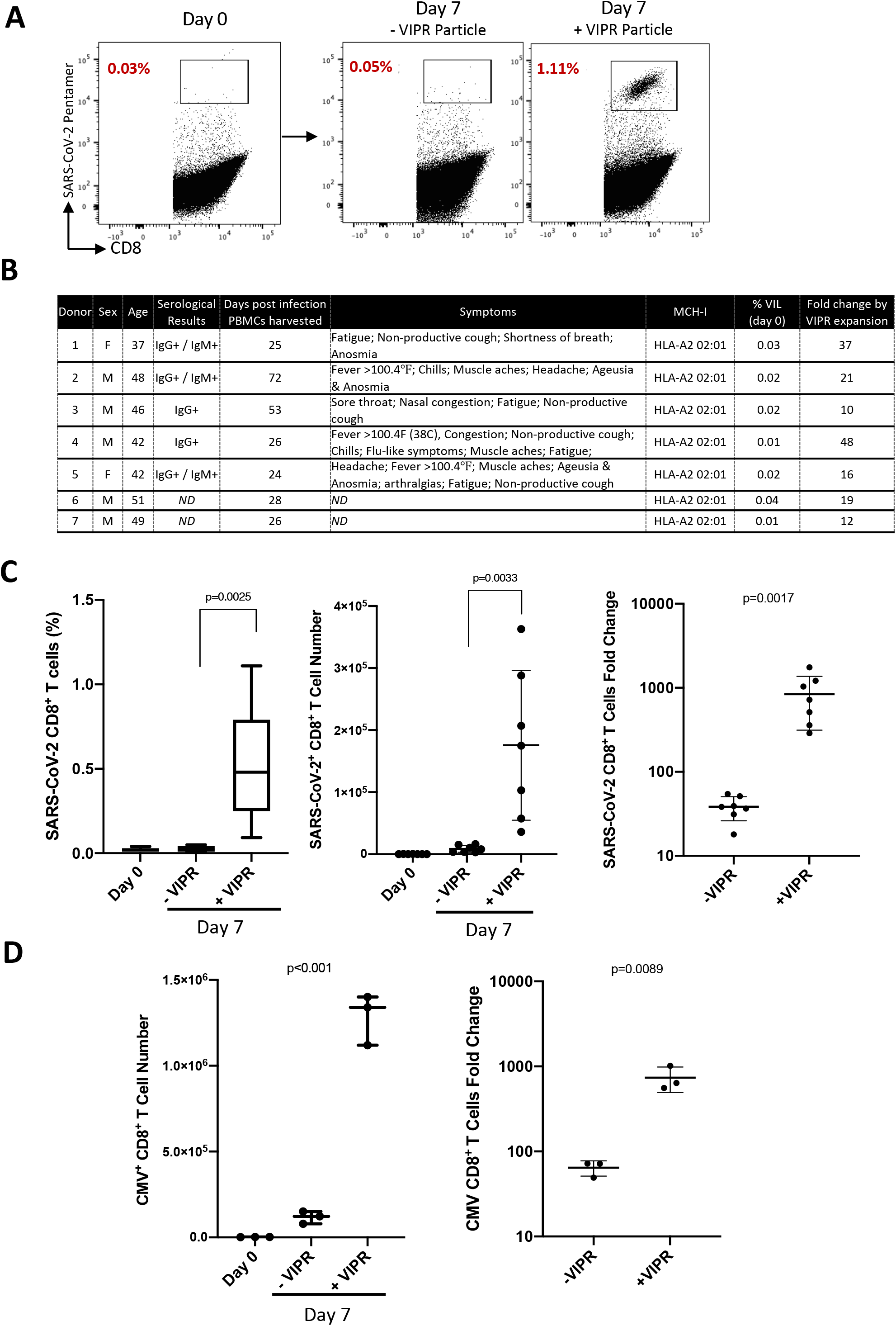
Rapid VIL expansion results in a 1,000-fold enrichment of SARS-CoV-2 antigen-specific T cells within 7-days from convalescent donors. **a,** Representative flow-cytometric analysis showing proportions of SARS-CoV-2 antigen-specific CD8^+^ T cells 7-days after expansion and enrichment with YLQPRTFLL antigen-specific VIPR Particles. **b**, Table describing the clinical presentation of convalescent COVID-19 donors used in this study and the proportions of SARS-CoV-2 specific VIL detectible before enrichment, and fold expansion by VIPR particles. **c**, Summary of CD8^+^ SARS-CoV-2 VIL %, cell number and fold change after VIPR expansion *(n=7).* **d**, Histograms showing total CMV pp65 antigen-specific CD8^+^ T cell numbers expanded by VIPR particles after 7-days and overall fold change in antigen specific cells *(n=3)*.

However, after 7-day culture with MHC-I VIPR particles, antigen-specific CD8^+^ T cells could be readily detected by pentamer staining and could be enriched and expanded to frequencies greater than 1% (Fig 2a&b). While the majority of PBMCs from convalescent individuals with the YLQ matched MHC allele (HLA A*02), included T cells from which antigen specific VIL could be enriched, the overall frequency varied between individuals and did not appear to correlate with either the length of time since they were symptomatic, nor the reported severity of their symptoms (Fig. 2b)

### Rapid VIL expansion results in 1,000-fold enrichment of CMV antigen-specific T cells within 7-days

Having demonstrated that VIPR particles can enrich both SARS-CoV-2 and CMV specific T cells, we evaluated the capacity for the 7-day culture system to expand the overall quantity of virus specific VIL. The T cell cultures were configured to include the same culturing conditions used in neoantigen TIL human clinical trials to enable rapid T cell expansion, thus in addition to the VIPR particles and high IL-2 (6000 IU/ml), IL-7, IL-15 and NAC, T cells were cultured in Gas Permeable Rapid Expansion (G-REX) plates. These culture plates enable gas exchange from the base of the culture well, allowing cells to be cultured with a large ratio of media per surface area and abundant access to nutrients, and have been shown to facilitate a large and rapid expansion of primary human T cells^32^.

After 7-days of culture with VIPR particles, SARS-CoV-2 antigen-specific CD8^+^ T cells could be robustly expanded in proportion, but most importantly in absolute quantity of T cells, to an average of over 1,000-fold (Fig. 2c). Thus, we found that cultures seeded with 1×10^6^ total CD3^+^ T cells could reach expanded numbers, on average, between 2.6×10^7^ and 4.5×10^7^ total cells at day-7. This proliferative expansion coupled with the enrichment of the VIL population resulted in an average of 2.4×10^5^ SARS-CoV-2 CD8^+^ T cells per million CD3^+^ cultured (Fig. 2c). A similar robustness in the expansion in absolute number of virus-specific CD8^+^ VIL could also be observed with T cells from CMV-positive individuals when stimulated with MHC-I VIPR particles under these culture conditions (Fig. 2d). After 7-days, antigen-specific CD8^+^ T cells had increased from approximately 2×10^3^ cells to over 1.0×10^6^, leading to up to an average >700-fold expansion in cell number.

Collectively these data demonstrate the ability of the VIPR particle expansion protocol to rapidly enrich and expand VIL from low numbers in CMV-positive individuals and near undetectable numbers in COVID-19 convalescent individuals, to significantly large numbers of virus-specific T cells.

### The activation and T cell memory phenotype of rapidly-expanded CMV and SARS-CoV-2 antigen-specific VIL

To evaluate the phenotype of SARS-CoV-2 specific T cells and CMV specific T cells that had undergone enrichment and expansion with VIPR particles, T cells were analyzed for expression of cell surface markers indictive of T cell activation. We observed SARS-CoV-2-specific CD8^+^ T cells expressing co-stimulatory and activation markers 4-1BB, OX-40 and CD25, albeit variable between convalescent individuals, and an elevated expression of HLA-DR when compared to non-virus-specific T cells within the culture (Fig. 3a). The SARS-CoV-2 antigen-specific VIL population also showed a significant expression of the checkpoint markers PD-1, TIGIT, LAG-3, indicating these T cells have acquired a proliferative and activated functional phenotype (Fig. 3b). The same profile of activation marker and checkpoint gene expression was observed when CMV-specific VIL were stimulated after 7-day rapid expansion with VIPR particles, with a similarly observed variability between different CMV-positive individuals, indicating this culture platform is effective at rapid T cells expansion and activation with multiple viral antigens (Fig. 3c&d).

**Fig. 3:**
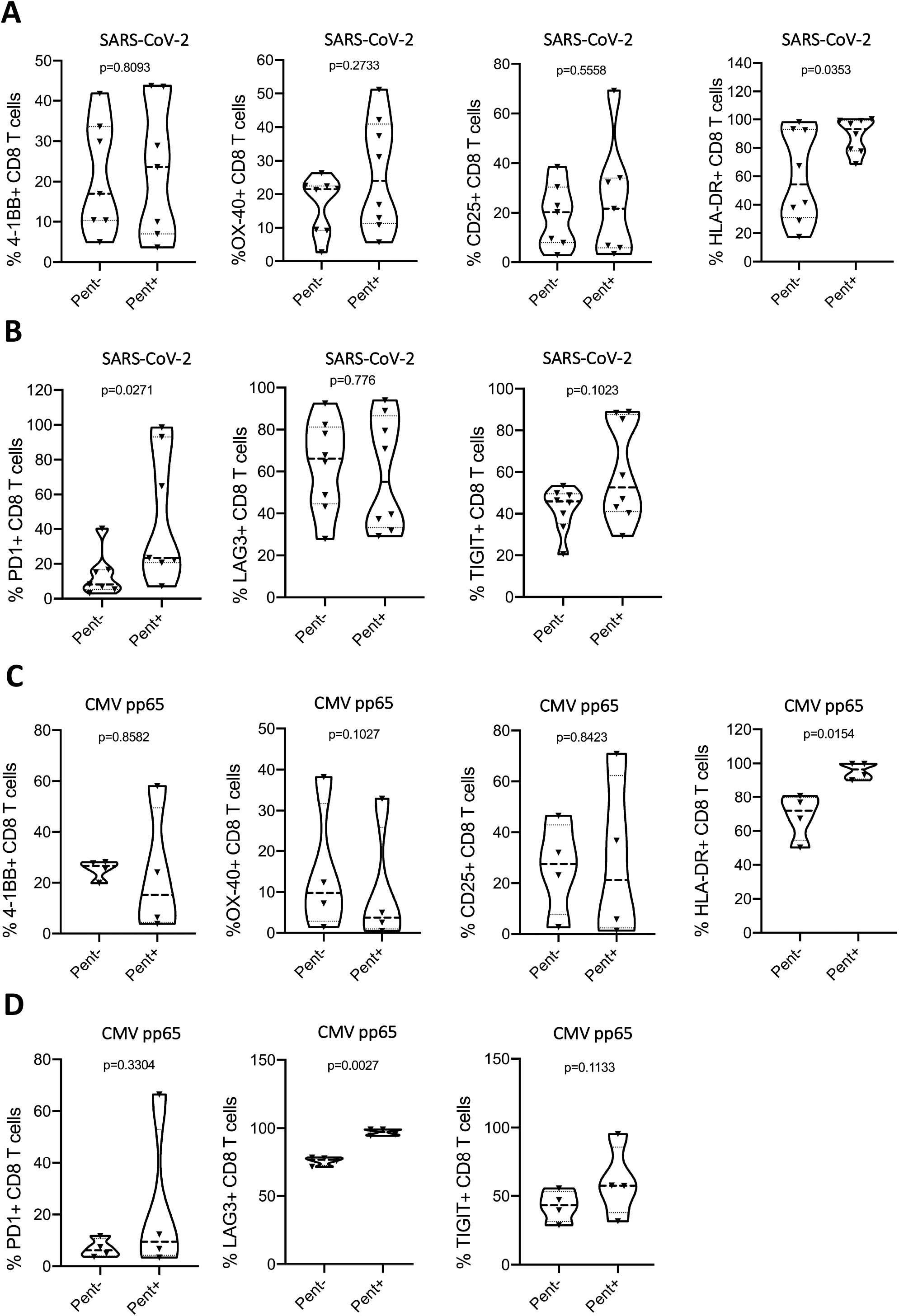
The activation and exhaustion phenotype of rapidly-expanded SARS-CoV-2 and CMV antigen-specific VIL. **a,** Expression of activation markers, 4-1BB, OX-40, CD25 and HLA-DR, and **b**, expression of checkpoint genes PD-1, LAG-3 and TIGIT among enriched and expanded SARS-CoV-2-specific CD8^+^ T cells at day 7 *(n=8)*. **c**, Analysis as in a, for CMV-specific CD8+ T cells and **d**, Analysis as in b, for CMV-specific CD8+ T cells *(n=4)*.

We analyzed the memory phenotype of the expanded SARS-CoV-2 and CMV virus-specific T cells by measuring expression of the canonical memory markers CD45RA and CD45RO and categorized the cell populations into either a naïve (CD45RO^−^, CD45RA^+^) or memory phenotype (CD45RO^+^, CD45RA^−^). After the 7-day culture in IL-2, IL-7, IL-15 and NAC, the majority of CMV T cells had begun to adopt a memory phenotype, but the virus-specific CD8^+^ T cells were almost exclusively expressing the highest levels of CD45RO and completely lost CD45RA expression, indicating the antigen-specific population had uniformly transitioned into memory T cells (Fig. 4a). Further delineation of the T cell memory phenotype by analysis of CD62L expression within the CD45RO^+^ population revealed the virus-specific T cells had robustly differentiated into an effector memory T cell phenotype via downregulation of CD62L (Fig. 4a&b). The non-virus-specific T cells within these cultures however, consisted of significantly more naïve T cells. The same profile of effector memory T cells was observed when SARS-CoV-2-specific VIL were stimulated after 7-day rapid expansion with VIPR particles, again demonstrating the antigen-specific VIPR particle platform is effective at significantly expanding activated effector memory T cells over a short time-course. (Fig. 4c&d).

**Fig. 4:**
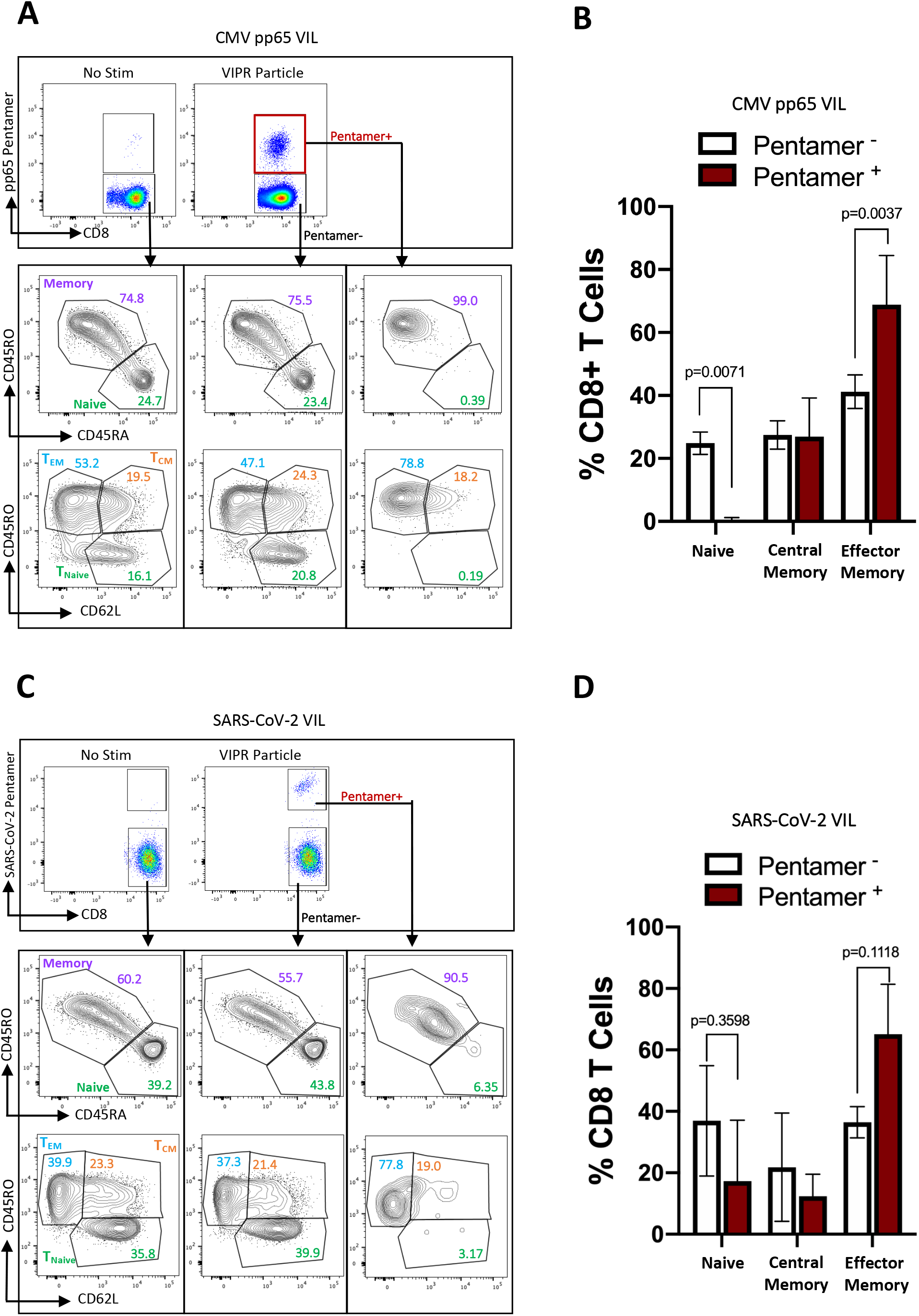
The T cell memory phenotype of rapidly-expanded CMV and SARS-CoV-2 antigen-specific VIL. **a,** Representative flow-cytometric analysis showing expression of CD45RO, CD45RA and CD62L among enriched and expanded CMV-specific CD8^+^ T cells at day-7. **b**, Summary of Naïve, Central memory and effector memory T cells subsets from data obtained in a, *(n=4)*. **c**, Analysis as in a, for SARS-CoV-2-specific CD8^+^ T cells, and **d**, analysis as in b, for SARS-CoV-2-specific CD8^+^ T cells *(n=8)*.

### Polyfunctional proinflammatory cytokine expression among rapidly-expanded CMV and SARS-CoV-2 antigen-specific VIL

To further evaluate the function of the rapidly expanded virus-specific VIL, we performed intracellular cytokine staining and flow cytometry and measured the proportion of the cells that were producing IFNγ, TNFα and IL-2. VIPR particle expanded T cells from convalescent COVID-19 individuals were stimulated after 7-day culture with 20μg/ml of SARS-CoV-2 YLQPRTFLL peptide antigen for 6-hours. We observed strong expression of all three proinflammatory cytokines within the antigen-specific T cell population (identified by TCR specific pentamer staining), but could not detect expression of either IFNγ or TNFα in the non-antigen-specific T cell population (T cells that do not bind the TCR-specific pentamer) nor within the T cells cultured for 7-days without any VIPR particle expansion (Fig. 5a&b). The antigen-specific T cells also showed significantly elevated levels of IL-2 when compared to the non-antigen specific CD8^+^ population. When analyzed together we see an elevated proportion of cells expressing 1, 2 or all 3 cytokines in combination when compared to the non-SARS-CoV-2-specific T cells within the expanded culture (Fig. 5c&d).

**Fig. 5:**
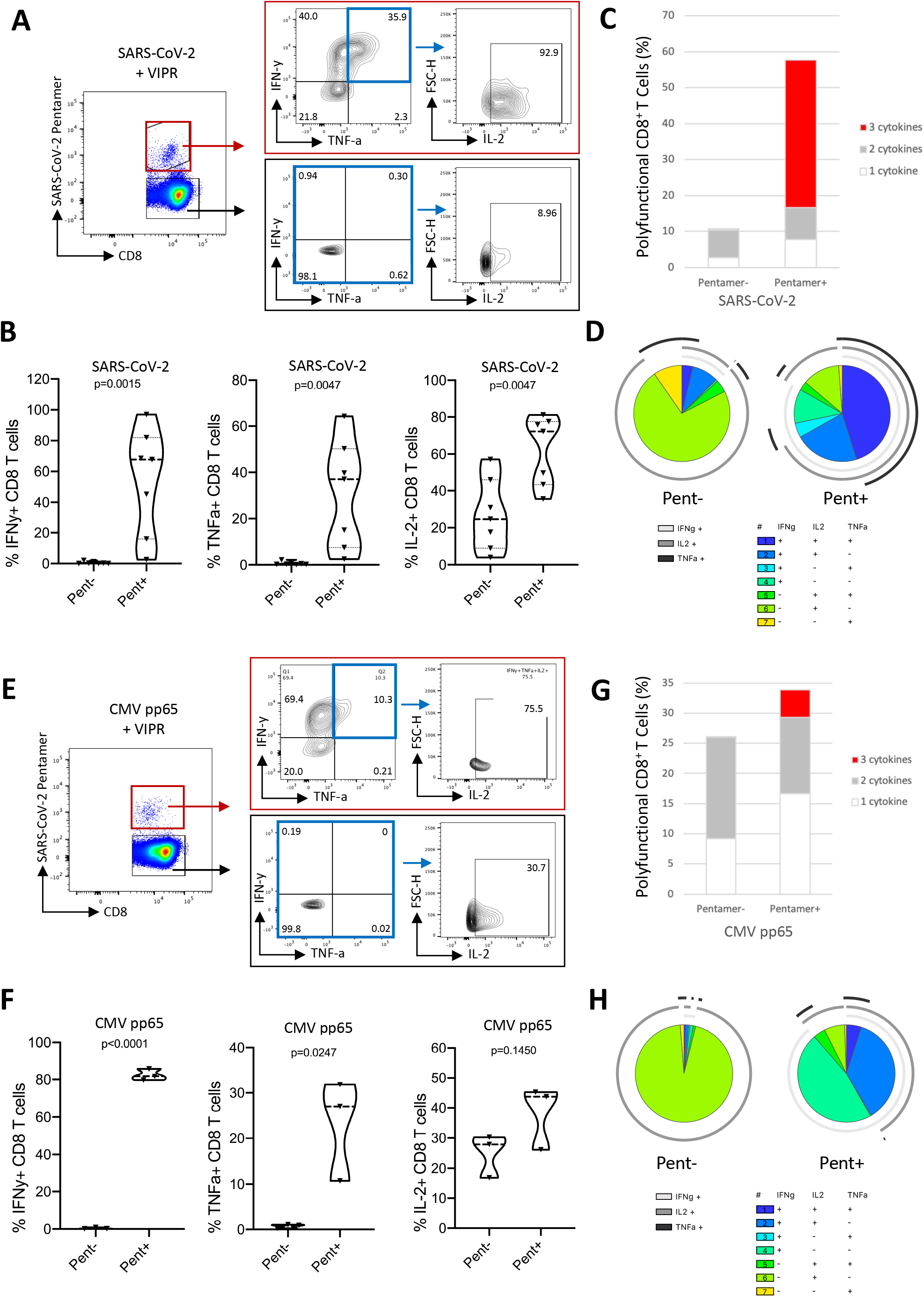
Proinflammatory cytokine expression among rapidly-expanded SARS-CoV-2 and CMV antigen-specific VIL. **a,** Representative flow-cytometric analysis showing expression of IFN-γ and TNFα among enriched and expanded SARS-COV-2 CD8^+^ T cells at day-7, after 6-hour stimulation with specific peptide antigen, and proportions of IFN-γ/TNFα expressing cells also expressing IL-2. **b**, Summary of the data obtained as in a for each cytokine *(n=7)*. **c**, Representative proportion of SARS-CoV-2 CD8^+^ T cells expressing 1, 2 or 3 cytokines. **d**, Extended analysis of SARS-CoV-2 VIL polycytokine function as SPICE representation. **e**, Analysis as in a, for CMV-specific CD8^+^ T cells. **f**, Histograms analyzed as in b summarizing the data obtained with CMV antigen specific CD8+ T cells *(n=3)*. **g**, Representative proportion of CMV CD8^+^ T cells expressing 1, 2 or 3 cytokines. **h**, Extended analysis of CMV-specific VIL polycytokine function as SPICE representation.

The same functional response was observed with virus-specific VIL expanded in T cells isolated from CMV-positive individuals and stimulated for 6-hours with pp65 MHC-I epitope peptide antigen. Intracellular cytokine staining revealed a robust increase in production of all cytokines in the CMV-specific CD8^+^ T cells when compared to the non-specific T cells from the same cultures or non-stimulated controls (Fig. 5e&f). An elevated frequency of polyfunctional CD8^+^ T cells expressing multiple proinflammatory cytokines was also seen in the CMV-specific T cell population (fig. 5g&h).

Taken together, these analyses demonstrate that elevated numbers of virus-specific VIL can be rapidly expanded in 7-days by VIPR particle culture and form robust activated, polyfunctional effector memory T cells.

## DISCUSSION

Given the paucity of therapeutic options for the treatment of COVID-19 and the provocative data suggesting the importance of the immune T cell response to viral infections, we investigate a novel potential therapeutic modality to augment the anti-viral T cell response by providing a therapeutic immune boost of virus-specific T cells (VIL). Similar to adoptive cell therapy (ACT) methods in immunooncology to actively transfer Tumor Infiltrating Lymphocytes (TIL), here we demonstrate the potential utility of Virus Induced Lymphocytes (VIL) to deliver SARS-CoV-2 immunodominant viral antigens in the setting of nascent and acute COVID-19 infection. Unlike T cell immuno-oncology in which T cell expansion and subsequent efficacy requires substantial critical mass of quantity, and thus time, VIL serve a catalytic immune booster function and can be isolated and expanded *ex vivo* both autologously and also MHC typed for allogenic delivery in a 7-day vein-to-vein time which is clinically practical and relevant (as depicted in fig. 6a).

**Fig. 6:**
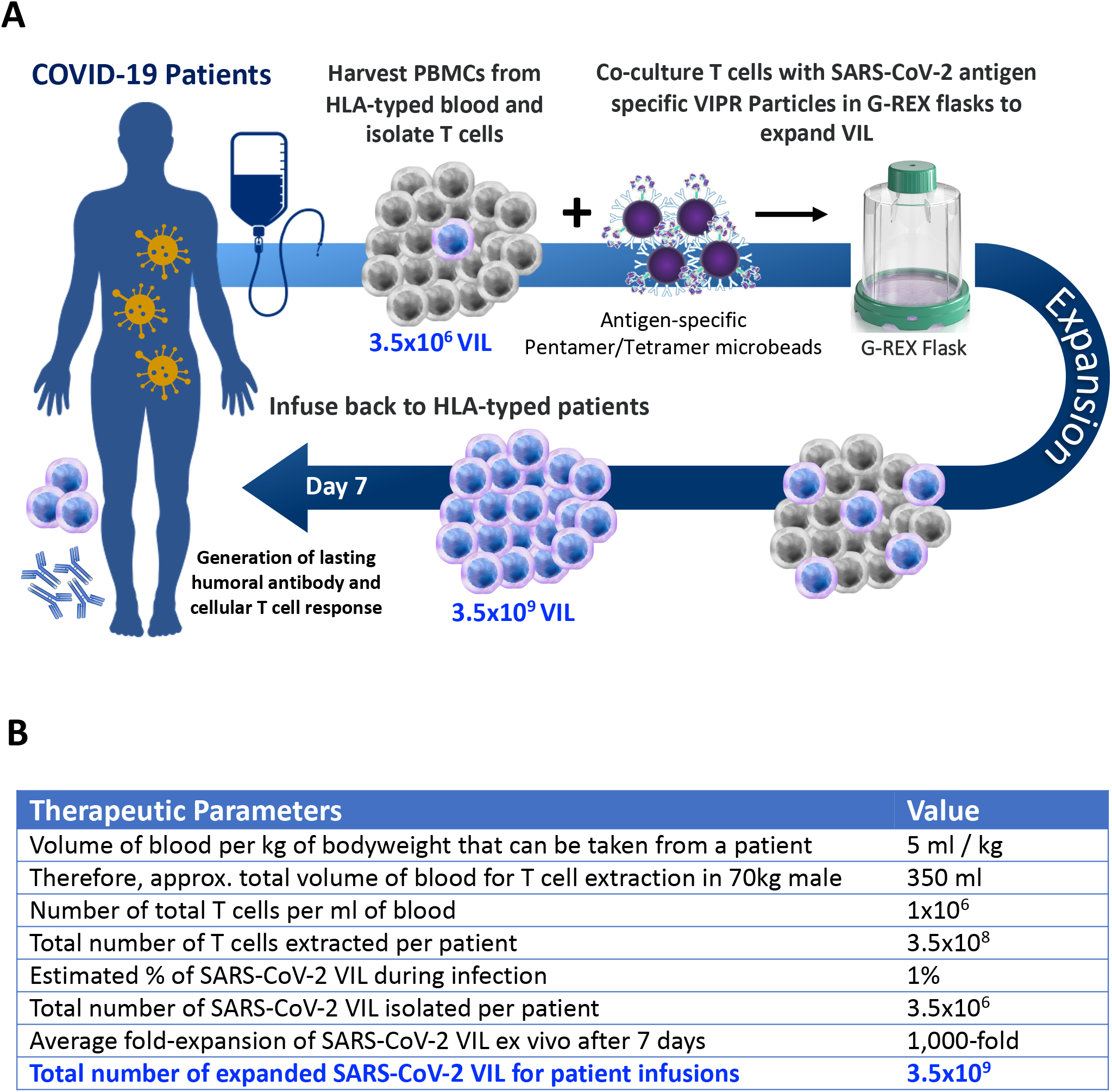
Allogeneic VIL Therapy Platform: An adoptive cell therapy for the treatment of individuals suffering from severe symptoms of COVID-19. **a,** Schematic for a COVID-19 cell therapy in which PMBCs are collected from the blood of HLA-typed hospitalized patients and total T cells isolated. T cells are stimulated with HLA-matched MHC-I/MHC-II antigen-specific SARS-CoV-2 VIPR beads to enrich and expand CD4^+^ and CD8^+^ T cells with TCRs specific for the SARS-CoV-2 antigen epitopes. These antigen-specific VIL expand at an average of 1,000-fold prior to adoptive transfer back to HLA-matched patients to mediate a T cell immune response to support the eradication of the SARS-CoV-2 virus and to engender protective immunity against repeat infection. **b**, Estimations of viral-specific T cell numbers generated ex vivo for patient infusion based on the empirical data of VIL expansion by VIPR particles.

In the setting of COVID-19 pathogenesis, studies have found individuals suffering from a more severe presentation of the disease typically require a duration of hospitalization ranging from 5 to 29 days^33, 34^. Thus, a therapeutic treatment to improve patient outcome must be rapidly administered during the critical time window of disease progression prior to and/or early in the patient’s intubation and ventilatory support. Herein lies the opportunity for a T cell therapy that can robustly expand the quantity and quality of virus-reactive T cells within this short duration to help boost the immune response and potentiate the patient’s own *in vivo* T cell response to the viral infection. Developments in Rapid Expansion Protocols (REP) for T cell and TIL expansion in the setting of clinical oncology have enabled methods for the robust and exponential *ex vivo* expansion of unenriched, as well as neoantigen-enriched, T cells for the autologous treatment of solid tumors^35, 36, 37, 38^. In fact, large quantities of antigen-specific TIL can be expanded in just 22 days using well established protocols^39^. This duration is however too long for a clinically relevant cell therapy in the setting of COVID-19 and thus in this study we have developed a more rapid T cell expansion protocol that builds upon the principles of TIL and T cell REP.

Our approach enables an over 1,000-fold expansion in the numbers of viral antigen-specific T cells in just 7-days from isolation of a patient’s T cells, providing a higher quantity of activated, polyfunctional effector memory cells. This study demonstrates the antigen-specific expansion of SARS-CoV-2 T cells from convalescent COVID-19 patients to demonstrate applicability for this viral pathogen, yet one important consideration is that these individuals have very low numbers of recirculating SARS-CoV-2 specific T cells in their blood due to the timeframe since their infection and recovery (fig. 2b). Hospitalized, symptomatic COVID-19 patients with a severe form of the disease will be undergoing a significant cellular immune response whereby the numbers of virus-specific T cells may have expanded, even if overall T cell numbers may be reduced in some individuals^40^. While this T cell response needs therapeutically boosting to potentiate viral clearance and ensure positive disease outcome, we expect a more robust and significantly elevated VIL expansion can be achieved when T cells are acquired from suffering COVID-19 patients as opposed to recovered convalescent individuals. When considering the translation of this platform into the clinic as a potential cell therapy for hospitalized COVID-19 patients, based on the level of enrichment and expansion demonstrated in this study, and a prediction of the number of SARS-COV-2 cells in the blood of COVID-19 individuals, we calculate an estimated capability to expand and deliver an average of approximately 3.5×10^9^ SARS-CoV-2 CD8^+^ and/or CD4^+^ T cells back to the patient within 7-days (Fig. 6b).

The key features of this VIL expansion platform include the use of high concentration of IL-2 to enable robust T cell proliferation supported by IL-7 and IL-15, and the addition of N-acetyl cysteine (NAC) which has been shown to significantly reduce upregulation of the DNA damage marker γH2AX, and the subsequent cell death seen in T cell culturing^41^. The expansion of VIL in G-REX plates and flasks also improves overall T cell proliferation by supporting more effective gaseous exchange and nutrient availability, further adding to the rapidity by which significant numbers of virus-specific VIL can be generated (Fig. 2c&d). A major component and innovation of the platform are the VIPR Particles themselves, which provide a viral epitope peptide in the context of MHC class I pentamers, or class II tetramers, to bind and stimulate the cognate TCRs specific for this antigen (fig. 1a). These aAPCs mimic the physiological presentation of viral antigens to CD8^+^ T cells by dendritic cells that occurs in local draining lymph nodes during viral infection, and the coating of the particles with anti-CD28 antibodies provides the necessary co-stimulatory signals that are critically required for effective T cell activation and formation of effector memory T cells^42^. It is well established that costimulatory signals like those delivered by CD28 (signal two) dictate whether CD8^+^ T cells will become optimally activated and expand, or whether the activation will be suboptimal^42^. While studies have also shown that T cells can indeed be expanded, from memory, and protect form viral infections in animal models in the absence of co-stimulation^43, 44^, it was found that very high levels of TCR stimulation were needed to overcome the need for co-stimulation^45, 46^. Thus, if a strong, shared, immunodominant and well validated peptide antigen is known and available for a viral pathogen, large numbers of VIL could potentially be expanded *ex vivo* by soluble peptide in the absence of an aAPC or co-stimulation. However, this situation is clinically irrelevant for most pathogens whereby multiple immunogenic epitopes typically exist and the comparative affinity for their cognate TCRs may not be known (as in an emerging pandemic). This is true in the case for the recently emerged SARS-CoV-2 virus which has been shown to harbor dozens of immunogenic shared epitopes spread throughout the genome, 90% of which are not located in the spike protein and show almost no cross-reactivity with known epitopes in seasonal coronaviruses, and for which T cell immunity must be established empirically^12, 31^.

Furthermore, a hospitalized COVID-19 patient’s own TCR repertoire for SARS-CoV-2 cannot be quickly assessed and determined to ensure they can respond to the strongest of these peptide epitopes in the absence of co-stimulation. Thus, VIL expansion platforms for newly emerged pathogens, as well as seasonal variants of existing viruses, will benefit from aAPC-mediated approaches, where the presence of CD28 co-stimulation should lower the T cell activation threshold and provide robust co-stimulation to ensure strong proliferation and effector memory T cell formation.

An alternative approach for VIL expansion by viral antigen presentation to T cells in the context of this co-stimulation, is the use of autologous professional antigen presentation cells (APCs) such as dendritic cells harvested from the patient’s blood, pulsed with antigenic peptide and co-cultured with the T cells. While in theory this works well for antigen-specific activation and expansion of CD8^+^ and CD4^+^ T cells and is routinely used in the setting of neoantigen-specific TIL expansion^38^, this is impractical in the setting of rapid VIL expansion as a therapeutic for hospitalized patients suffering from acute viral infections such as with SARS-CoV-2. Dendritic cells are typically generated from autologous monocytes through a series of *ex vivo* maturation steps with different cytokines which includes approximately 6 days of culture in IL-4 and GM-CSF to generate immature DCs before further maturation^47^. This requirement for several days of manipulation before the dendritic cells can be used to activate and expand the patient’s VIL, precludes their use for rapid T cell expansion in the setting of acute and severe viral infections such as with COVID-19. Therefore, we believe the VIPR platform based on artificial APCs offers a clinically meaningful and practical alternative to this approach.

Given the importance of the aAPC VIPR particles for T cell stimulation, we optimized the design to maximize the strength of the TCR and CD28 engagement by determining both the effective dose ratio of particles per T cell (1:20) and the optimal density of peptide-MHC-pentamer versus anti-CD28 antibody (1:30) on the surface of the particle (fig. 1d&e). When the optimized VIPR particles were combined with other elements of the protocol, including high cytokine concentrations and G-REX flasks, we thus observed not only a significant expansion in virus-specific VIL cell numbers (Fig. 2c&d), but that these cells had developed into activated effector memory T cells (Figs. 3&4). This was crucial to establish a pool of virus-reactive T cells that can not only mount a primary immune response against the virus when adoptively transferred to the patient, but also engender immunological memory to ensure the duration of the response is long lasting. Given the active state of the VIL by day-7, it is likely that further cellular expansion will be catalyzed upon ACT to the patient given the characteristic upregulated endogenous inflammasome of COVID-19 patients. Importantly, after a 7-day expansion of the SARS-CoV-2 VIL, the cells were producing robust levels of polyfunctional cytokine (Fig. 5). Thus, the expanding virus-specific VIL that can subsequently be transferred back to the patient are primed to mount a functional cellular response in the context of cytokine release and recruitment of other immune cells. These antigen-specific CD8^+^ T cells are also likely primed to recognize and mount a cytolytic response against infected cells, aiding to clear the viral infection.

The VIPR rapid expansion platform is modular and tunable to multiple viral antigens restricted to different MHC alleles. Using validated CMV epitopes, we generated MHC class II VIPR particles against the DRB1*07:01 restricted gB 215-229 antigen and were able to enrich and expand CD4^+^ T cells (Fig. 1b). Thus, this technology can be used to expand both T Helper and Cytolytic virus-specific VIL to provide a modality to tune a specific cell therapy product to treat different viral diseases. The enrichment and expansion of CMV-specific T cells demonstrates the flexibility of the VIPR particle system for addressing cell therapies for diverse pathogens where immunogenic epitopes are known. VIPR particles are off-the-shelf reagents that can be rapidly utilized for *ex vivo* T cell expansion in the clinic without any additional manufacturing lead time. We expect that this technology would provide a modality for expansion of all clinically relevant virally-mediated diseases where an immune boosting dose of virus-specific T cells would aid in viral clearance and disease outcome. Furthermore, due to the immune catalytic nature of T cells and their ability to expand further *in vivo* and recruit additional cells of the immune system, we believe that an efficacious therapeutic dose of virus-specific T is likely to be well within the numbers achievable within short durations of expansion. In instances whereby such a short duration of T cell expansion is not as critical as required in diseases like COVID-19, this technology can also enable the expansion and cryopreservation of virus-specific VIL.

To be broadly applicable as a pragmatic modality that is scalable, we have MHC typed donor pools of SARS-CoV-2-specific VIL for use as an allogeneic off-the-shelf therapy at scale. In this setting, these T cells are tissue matched to COVID-19 patients and provided as an allogeneic cell therapy product to combat an active infection. Thus, by virtue of having pre-expanded stocks of SARS-COV-2 cells from SARS-CoV-2 donors, hospitalized patients can be treated immediately with the cells within the ICU upon MHC tissue type results. Furthermore, cryopreservation of other VIL, such as MHC - typed CMV-specific VIL, could, as an example, be used to immunize bone marrow transplant recipients or other immunocompromised individuals against adventitious pathogens^48^.

In summary we demonstrate a novel technology platform for the robust and rapid expansion of virus-specific T cells as a potential cell therapy for COVID-19 and other viral pathogens. Heretofore, the promising antigen-specific therapeutic approaches to date, including polyclonal antibody cocktails and monoclonal antibodies, and current prophylactic vaccine approaches to COVID-19, all have been focused on neutralizing antibodies. However, complete immune protection is likely also a function of the long-lasting central memory T cell response to provide both cellular immunity, and potentiate humoral immunity, and thus prolong the duration of protection^8, 11^.

As development and validation of the VIL platform continues, a global “second-wave” of COVID-19 morbidity and mortality, perhaps exacerbated by seasonal inflection, appears to be accelerating^49^. Validated therapies are few and largely supportive or non-specific, such as the use of dexamethasone to delay mechanical ventilation in COVID-19 induced pulmonary failure. The morbidity and mortality of COVID-19 critical illness remain alarmingly high^50^. Thus, there is an urgent need for specificity and efficacy in the clinic to mitigate disease progression and severity. Using the same cGMP validation processes employed in the setting of a CRISPR-engineered TIL cell therapy for cancer, we will explore a timely regulatory path to evaluating the VIL platform technology in a phase I/II clinical trial of SARS-CoV-2-infected patients, should there continue to be an urgent need during the course of the COVID-19 pandemic.

## METHODS

### Patient samples and preparation of T cells

Peripheral blood mononuclear cells were obtained from anonymized CMV-positive individuals and convalescent COVID-19 individuals (Caltag Medsystems, Tissue Solutions Ltd, Precision for Medicine, Inc.) and obtained, handled and stored in accordance with the Human Tissue Authority UK regulations. Genomic DNA was extracted from PBMC samples using the Gentra Puregene DNA isolation kit (Qiagen) and DNA samples were HLA-typed by sequencing at Class I (HLA-A, -B & -C) and Class II (HLA-DRB1) loci (MC Diagnostics). Total CD3^+^ T cells were isolated from unfractionated PBMCs using the EasySep™ Human T Cell Isolation Kit (Stem Cell Technologies) with a DynaMag™-2 magnet (ThermoFisher Scientific) according to the manufacturer’s guidelines. Purity and viability of isolated T cells was assessed using flow cytometry prior to cryopreservation of isolated T cells in CryoStore CS10 cryopreservation media (Stem Cell Technologies) at a density of 1-1.5×10^7^ cells per ml.

### Preparation of VIPR Particles

Micro-aAPC VIPR particles were constructed by direct conjugation of biotin labelled peptide-MHC-Pentamers and biotinylated MHC class II Tetramers to streptavidin Dynabeads in combination with biotinylated anti-CD28 antibodies. Pro5 MHC Class I Pentamers were provided by ProImmune Ltd and include the following peptide epitopes: CMV pp65 HLA A*02:01-restricted NLVPMVATV epitope; SARS-CoV-2 Spike protein 269-277 HLA A*02:01-restricted YLQPRTFLL epitope. ProM2 MHC class II biotinylated Monomers were also obtained from ProImmune for the DRB1*07:01-restricted CMV gB 215-229 PDDYSNTHSTRYVTV epitope. These biotin-labelled MHC-peptide Pentamers and biotinylated MHC-peptide Monomer complexes, and mouse anti-human CD28 antibody (BD Biosciences) were conjugated to M270 Streptavidin Dynabeads (Thermo Scientific) at defined molar ratios of Pentamer:anti-CD28 and Monomer:anti-CD28 (calculated to account for tetramer formation of these monomers), and both in molar excess of the number of streptavidin molecules per Dynabead. Bead-conjugation was carried out at 4°C in Phospho-buffered saline (PBS, Gibco), for 1 hour with regular agitation. Conjugated VIPR particles were centrifuged at 14,500 xg for 3 minutes at 4°C and washed three times in PBS while the beads were immobilized in an Invitrogen DynaMag-2 magnet (Thermo Scientific). VIPR particles were resuspended in sterile PBS at a concentration of 5×108 particles per ml and stored at 4°C.

### Expansion of antigen-specific T cells by VIPR particles

Isolated human CD3^+^ T cells were cultured in X-VIVO-15 Basal Media (Lonza) supplemented with 10% Human AB Serum Heat Inactivated (Sigma), 6000IU/ml Recombinant Human IL-2 (Peprotech), 5ng/ml Recombinant Human IL-7 (Peprotech), 5ng/ml Recombinant Human IL-15 (Peprotech) and 10mM N-Acetyl-L-cysteine (Sigma). T cells were seeded at a density of 2×10^5^ cells per well of U-bottom 96-well plates, or at a density of 1-2×10^6^ T cells per cm^2^ of G-REX 24-well plates (Wilson-Wolf). At the time of T cell seeding, VIPR particles were added to the relevant samples at a ratio of 20 T cells per particle, and cells were cultured for 7-days in a 37°C, 5% CO2 culture incubator. In addition, a sample of the T cells was also analyzed by flow cytometry at Day-0 to measure the starting proportion of antigen-specific T cells (see methods below). For some cultures the media was replaced every 2-3 days with fresh complete media including cytokines and NAC (but no extra addition of VIPR particles) and media in G-REX cultures was left unchanged for the duration of 7-days in some experiments to promote cell expansion. On day 7, T cell numbers were assessed by harvesting all cells, washing in PBS followed by centrifugation at 300 xg for 10 minutes and then counting using a CellDrop Automated Cell Counter (DeNovix). The proportion of expanded antigen-specific T cells was assessed at Day-7 by flow cytometry.

### Flow cytometry analysis of T cell phenotype

For flow cytometric analysis of the antigen-specific T cell population and cell surface marker expression, cells were harvested from culture plates and washed using PBS with 1% Bovine Serum Albumen (Thermo Scientific) and were then stained with monoclonal antibodies specific for CD8 (HIT8A, 1:100), CD4 (OKT4, 1:100), HLA-DR (L243 1:80), LAG-3 (11C3C65, 1:80), TIGIT (VSTM3, 1:40), CD45RO (UCHL1, 1:40), CD45RA (HI100, 1:80), TIM3 (F38-2E2, 1:40), CD62L (DREG-56, 1:40), CD57 (QA17A04, 1:80), PD-1 (EH12.1, 1:40), OX-40 (Ber-ACT35, 1:40), CD25 (MA2-51, 1:40), 41BB (4B4-1, 1:40), (Biolegend) or specific for CD8 (RPA-T8, 1:100) (BD Bioscience), or TNF-a (MAb11, 1:40) and CD3 (UCHT1, 1:100) (ThermoFisher). CMV pp65 and SARS-CoV-2 Spike antigen-specific T cells were detected by staining cells with R-PE-labelled Pro5 Pentamers (ProImmune, Ltd), and CMV gB215-229 specific T cells stained with R-PE labelled ProT2 Tetramer for 20 minutes at room temperature according to manufacturer’s recommendation. Live/Dead Fixable Dead Cell Stains (Invitrogen) or SYTOX Blue Dead Cell Stain (Invitrogen) were included in all experiments to exclude dead cells. After staining, cells were resuspended in PBS with 2% Human Heat Inactivated AB Serum (Sigma) and 0.1M EDTA pH 8.0 (Invitrogen) before analysis on a Fortessa flow cytometer (BD Bioscience) and data analyzed using FlowJo 10 software (BD Biosciences).

### Intracellular cytokine staining

After 7-days of expansion with VIPR particles, the T cell cultures were stimulated for 6 hours with 20μg/ml peptide antigen (>95% purity) specific for the VIPR particle expanded CD8^+^ population (CMV pp65: NLVPMVATV; SARS-CoV-2 S protein 269-277 YLQPRTFLL), all peptides were synthesized and obtained from ProImmune, Ltd. After 1-hour of peptide stimulation, GolgiStop solution was added (containing Monensin protein transport inhibitor) to block intracellular protein transport (BD Bioscience). As a positive control for cytokine production, T cells were also stimulated for 6 hours with 50ng/ml PMA and 1μg/ml Ionomycin (Sigma). T cells were then harvested, and cells fixed and permeabilized using BD Cytofix/Cytoperm Fixation/Permeabilization Solution (ThermoFisher). Cells were then stained for surface markers followed by intracellular cytokines using antibodies specific for IFN-y (4S.B3, 1:40) (Biolegend) IL-2 (MQ1-17H12, 1:40) (BD Bioscience), or TNF-a (MAb11, 1:40) (ThermoFisher). Flow analysis was carried out on a Fortessa flow cytometer (BD Bioscience) and data analyzed using FlowJo 10 software (BD Biosciences).

### Statistical analyses

Statistical differences between two sample groups, where appropriate, was analyzed by a standard Student’s two-tailed, non-paired, t test using GraphPad Prism Software version 8. P values are included in the figures where statistical analyses have been carried out.

## Ethics declarations

D.B. has a sponsored research collaboration funded by Intima Bioscience. Intima Bioscience has patents filed based on the findings described herein. The authors declare no competing interests.

